# Syndecan-1 is overexpressed in human thoracic aneurysm but is dispensable for the disease progression in vivo

**DOI:** 10.1101/2021.12.16.471096

**Authors:** Sara Zalghout, Sophie Vo, Véronique Arocas, Soumaya Jadoui, Eva Hamade, Bassam Badran, Olivier Oudar, Nathalie Charnaux, Yacine Boulaftali, Marie-Christine Bouton, Benjamin Richard

## Abstract

Glycosaminoglycans (GAGs) pooling has been considered since long as one of the histopathological characteristics defining thoracic aortic aneurysm (TAA) together with smooth muscle cells (SMCs) apoptosis and elastin fibers degradation. However, few information is provided about GAGs composition or potential implication in TAA pathology. Syndecan-1 (Sdc-1) is a heparan sulfate proteoglycan that is implicated in extracellular matrix (ECM) interaction and assembly, regulation of SMCs phenotype and various aspects of inflammation in the vascular wall. In the current work, the regulation of Sdc-1 protein was examined in human TAA by ELISA and immunohistochemistry. In addition, the role of Sdc-1 was evaluated in descending TAA in vivo using a mouse model combining both aortic wall weakening and hypertension. Our results showed that Sdc-1 protein is over expressed in human TAA aortas compared to healthy counterparts and that SMCs are the major cell type expressing Sdc-1. Similarly, in the mouse model used, Sdc-1 expression was increased in TAA aortas compared to healthy samples. Although its protective role against abdominal aneurysm has been reported, we observed that Sdc-1 was dispensable for TAA prevalence or rupture. In addition, Sdc-1 deficiency did not alter the extent of aortic wall dilatation, elastin degradation, collagen deposition, or leukocyte recruitment in our TAA model. These findings suggest that Sdc-1 could be a biomarker revealing TAA pathology. Future investigations could uncover the underlying mechanisms leading to Sdc-1 expression alteration in TAA.

## INTRODUCTION

The mortality rate due to thoracic aortic aneurysm (TAA) rupture is in considerable increase and the efficiency of current drug therapies is yet limited and controversial (1–3). Glycosaminoglycans (GAGs) pooling has been considered since long as one of the histopathological characteristics defining TAAs, together with smooth muscle cells (SMCs) apoptosis or loss of contractility and elastin fibers degradation of (4–6). However, the identity of the accumulated GAGs in TAA and their potential involvement in the disease are poorly studied. A better understanding of the molecular changes occurring in TAA pathophysiology may open up new avenues for its treatment.

Even though proteoglycans (PGs; i.e. core protein with attached GAGs) occupy 1 to 5% mass fraction of a healthy arterial wall, they are crucial contributors to the aortic structural integrity and functioning and in particular to its mechanical homeostasis (7). PGs accumulation may exert a swelling pressure on the elastic laminae resulting in their separation and a consequent modification of the wall mechanical properties giving rise to a weakened wall and participating in TAA dissections and ruptures (7,8). Recently, it has been reported that aggrecan and versican, two chondroitin sulfate PGs, accumulate in regions of medial degeneration in ascending TAAs and dissections and may contribute to extracellular matrix (ECM) disruption (9). In addition, the levels of heparan and chondroitin sulfate PGs are increased after vascular injury and in dissected aortas (10–12).

Syndecan-1 (Sdc-1), a transmembrane heparan sulfate PG, is involved in various aspects of inflammatory diseases (13–15), wound healing (16), and ECM interaction or assembly (17,18). Sdc-1 has been shown to maintain a differentiated (19) and a contractile state of SMCs (12). In addition, it mediates cell-cell and cell-matrix interactions by acting as a co-receptor for binding to ECM molecules or growth factors such as fibronectin or VEGF, respectively (20,21). Therefore, Sdc-1 is hypothesized to play a role in the development of aortic wall pathologies. Sdc-1 has indeed been shown, in two mice models of abdominal aortic aneurysm (AAA) in which aneurysm was induced by elastase or angiotensin II (Ang II) infusion (on Apo E deficient background) (22), to exert a protective role against AAA development by attenuating the inflammatory response and reducing protease activity (22). In contrast, no data are available regarding its role in TAA development.

In the current study, we investigated Sdc-1 expression in the aortic wall of patients with TAA arising from different etiologies. We also examined the disease progression and characteristics by the use of an aneurysm mouse model generated by the combination of aortic wall weakness and hypertension in Sdc-1^+/+^ or Sdc-1^-/-^ on C57Bl/6J background.

## MATERIALS AND METHODS

### Human samples

Twenty-five human ascending TAA samples were obtained from patients at the time of prophylactic surgical repair and eleven healthy thoracic aortic tissues were obtained from anonymous deceased organ donors under the authorization of the French Biomedicine Agency (PFS 09-007). Data relative to these patients are stated in Supplementary Table 1. Aortic preparation involved direct paraformaldehyde fixation for histological studies or immediate macroscopic dissection to separate the distinct aortic layers (intima, media, adventitia) followed by direct freezing for q-PCR and ELISA analysis.

**Table 1:**
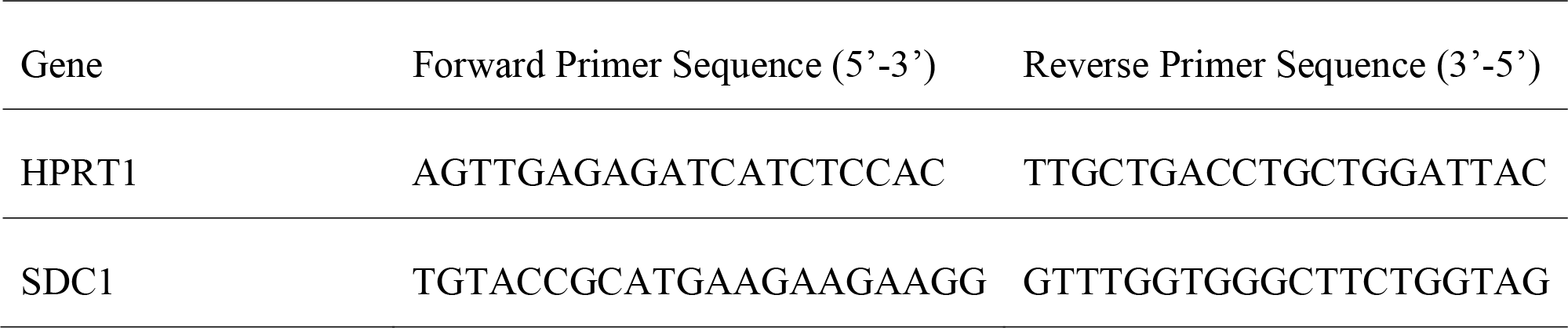
Primers used for quantitative polymerase chain reaction (qPCR; human samples)

### Quantification of mRNA and protein levels of Sdc-1 in human samples

Frozen media tissues from healthy or TAA aortas were pulverized using liquid nitrogen freezer mill (6870 SPEX Certiprep 6750). Protein extraction was done by powder lysis in RIPA buffer (5mg/ml). Sdc-1 media protein level was assessed using ELISA (R & D systems, DY2780) according to the manufacturer’s instructions. For RNA extraction, 50 to 100 mg of powder were homogenized in Trizol Reagent (Invitrogen). Total RNA was extracted using RNeasy extraction kit (Qiagen) and 1 µg was reverse transcribed with RT Maxima First Strand kit (K1642, Thermo Scientific). The q-PCR was performed on cDNA using Light Cycler (Roche). Levels of mRNA were normalized to hypoxanthine guanine phosphoribosyl transferase (HPRT). Sequences of primers are listed in table 1. Fold changes of gene expression were calculated using ΔΔCT method.

### Histology and immunostaining

For histological studies, human and murine fixed aortic tissues were embedded in paraffin, and sectioned into 7 μm thick sections. Antigen retrieval was done by heating the sections in a water bath at 95°C for 30 min in a Dako Target Retrieval Solution, pH 9 (code S236). Sections were permeabilized by 0.1% Triton-X 100 for 5 min. For DAB IHC endogenous peroxidase activity was blocked by 3% H_2_O_2_ for 30 min. Blocking was performed with DAKO^®^ protein block-serum free (code # X0909) for 1 h at room temperature. Sections were incubated overnight at 4°C with primary antibodies (Supplementary Table 2).

For DAB IHC, slides were washed with PBS and treated 1 h at room temperature with LSAB2 System, HRP kit for human, and Peroxidase AffiniPure donkey anti-rat secondary antibody for mice samples (Supplementary Table 2). 3′-diaminobenzidine tetra-hydrochloride chromogen (DAB, K3468, Dako) was added to all sections and the reaction was stopped with distilled water. Counterstaining was done with hematoxylin (for mice IHC). Slides were then mounted with Eukitt® mounting media.

For immunofluorescence, following primary antibody incubation, sections were treated with adequate secondary antibodies (Supplementary Table 2) for 2 h at room temperature. Nuclei were then stained with DAPI aqueous fluoroshield mounting media (Abcam, ab104139). α-SMA, CD45 and ly6-G markers were used to identify SMCs, leukocytes and neutrophils, respectively.

Elastin laminae degradation of mice aortas was blind-scored after orcein (Sigma-Aldrich) staining, using an ascending scale from 0 to 4 (no degradation to maximal degradation). Collagen deposition was detected with Sirius red (RAL Diagnostics) staining, and its quantification was assessed by image J software after exposing the slides to polarized light (Leica microsystems, DMi8).

### Animals and experimental procedures

Sdc-1^-/-^ mice were a kind gift from Dr. Pyong Woo Park (Harvard Medical School, Boston) and have been backcrossed for at least 10 generations on a C57BL/6J background.

Three week old Sdc-1^+/+^ and Sdc-1^-/-^ mice were divided into three groups: (group 1) control group with no treatment (mice were sacrificed at 8 weeks of age), (groups 2 & 3) treated groups including mice receiving daily intraperitoneal injection of 150 mg/kg/day of β-aminopropionitrile fumarate (BAPN, A3134; Sigma Aldrich) for 28 days, then infused subcutaneously with 1 µg/kg/min of angiotensin II (Ang II, A9525, Sigma Aldrich) during 3 (group 2) or 28 days (group 3) using mini-osmotic pumps (Alzet 2004) (Figure 2A). At the end of the protocol, mice were anesthetized and sacrificed. Aortas were harvested and rinsed with saline to remove blood. Aortas were cleaned from the surrounding connective and adipose tissues, fixed in 10% formalin for 24 h at 4°C, and stored in 70% ethanol for further investigations. Necropsy was done for mice that died during the course of the experiment and the site of rupture (thoracic or abdominal) was determined based on macroscopic view of hemorrhage location. Aneurysm (TAA or AAA) was defined as 50% increase in the mean aortic diameter compared to the same healthy aortic segment from non-treated mice (please check below quantification section).

### Blood pressure measurement

Mice blood pressure was measured non-invasively before pump implantation and up to 5 days post-implantation, using the tail cuff system (BP-2000 SERIES II, Blood Pressure Analysis System ™, Visitech Systems). Mice were habituated for a minimum of 5 consecutive days before the recording were considered.

### Measurement of aortic diameter

For morphometric analysis, images were taken by an EF-S 60 mm macro lens mounted on a DSLR camera (Canon EOS600D), and used to measure the outer diameter of the early descending thoracic aorta using Image J software. The diameter was determined at the site of dilatation from the average of a minimum of 3 different measurements of the posterior and inferior sides.

### Statistical analysis

Values are shown as percentage or mean ± standard error of the mean (SEM). Statistical analysis was performed using Prism GraphPad. The non-parametric Mann-Whitney test was used to compare two groups when data did not display a normal distribution. Fischer’s exact test was used when comparing two categorical variables. P values less than 0.05 were considered significant.

## RESULTS

### Sdc-1 protein expression is increased in the aortic media of patients with TAA and is expressed by SMCs

The expression of Sdc-1 at the mRNA and protein levels was investigated in human healthy or TAA medial layer. Similar levels of *SDC1* mRNA were observed between the different samples (Figure 1A). In contrast, a significantly higher protein level of Sdc-1 was revealed by ELISA in TAA compared with healthy human aortic media (Figure 1B). This difference was confirmed by IHC (Figure 1C). Immunofluorescence results showed a co-localization between Sdc-1 and α-SMA, indicating that most of Sdc-1 is expressed by SMCs in human healthy and TAA aortas (Figure 1D).

**Figure 1.**
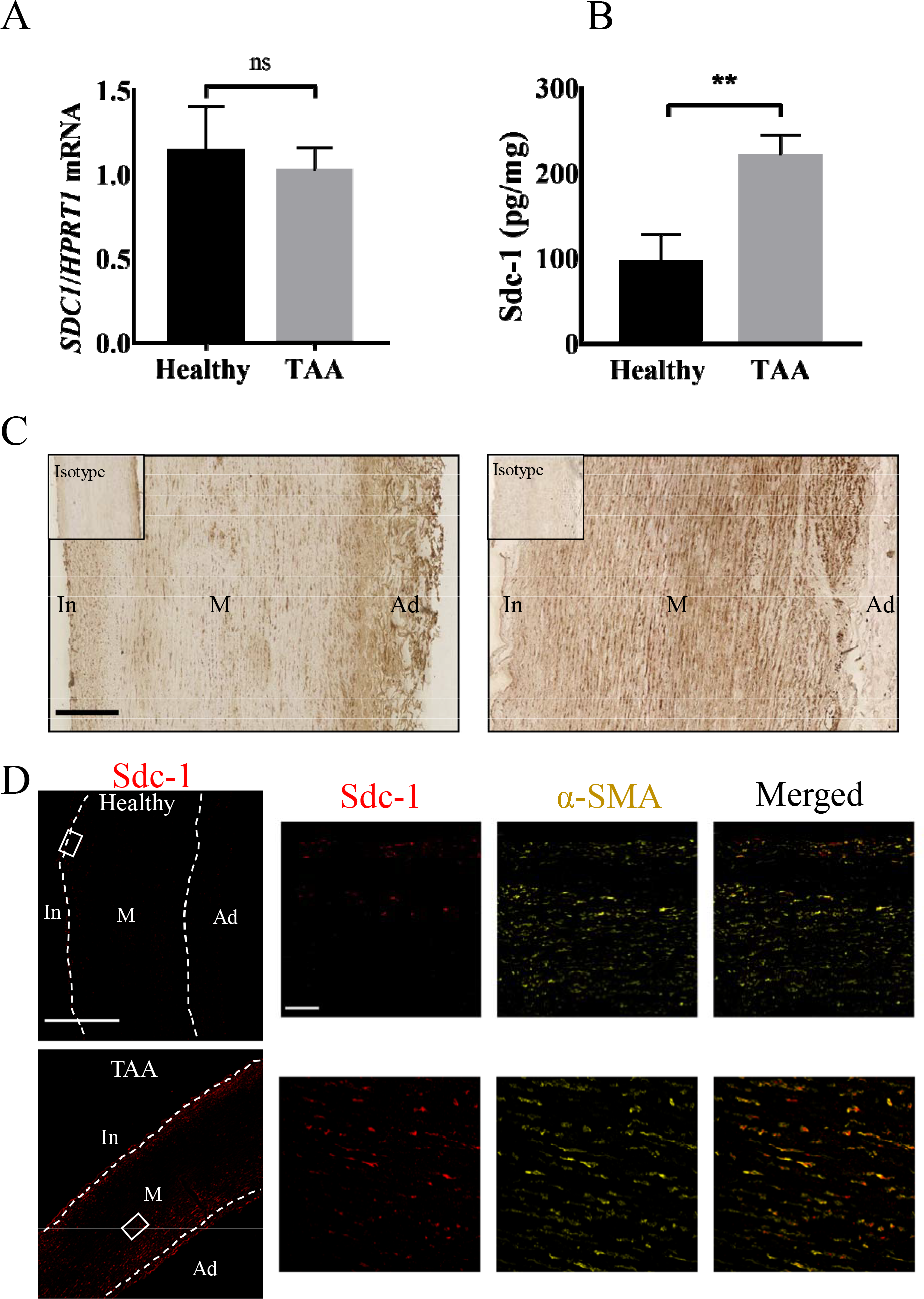
Sdc-1 protein level is increased in human TAA media compared to healthy ones and is expressed by SMCs. **(A)** mRNA level of *SDC1* was measured by q-PCR and normalized to Hypoxanthine-guanine phosphoribosyltransferase (HPRT) in healthy *(n=6)* and TAA *(n=14)* human thoracic aortic media. **(B)** The concentration of Sdc-1 protein in human aortic media was assessed by ELISA from healthy *(n=8)* or TAA *(n=16)* donors. **(C)** Representative images of Sdc-1 staining by immunohistochemistry in human healthy and TAA aortas. Staining with an isotype control was performed as a negative control and shown on the top left of images. Scale bar corresponds to 200µm. **(D)** Representative immunofluorescence images of Sdc-1 (red) and α-SMA (yellow) co-staining in human healthy (top images) or TAA (bottom images) aortas. The dashed lines indicate the separation of the different layers: In: intima, M: media, Ad: adventitia. Scale bar corresponds to 400µm for the non-magnified images and 50µm to the magnified ones. **(A, B)** Data are presented as mean ± SEM and *P* values were calculated using two-tailed Mann-Whitney test;** corresponds to *p value* <0.01, ns: not significant.

Altogether, these data illustrate that Sdc-1 is more expressed at the protein level in TAA aortas compared to healthy counterparts and SMCs are a type of cells overexpressing Sdc-1 in TAA aortas.

### Sdc-1 is overexpressed in the TAA developed in the BAPN/Ang II aneurysm mouse model

The role of Sdc-1 on TAA development was investigated in an animal model using BAPN and Ang II known to induce aortic aneurysms in C57Bl/6J mice (23). In the present model, 3-weeks-old Sdc-1^+/+^ or Sdc-1^-/-^ male mice received intraperitoneal injection of BAPN for 4 weeks and then subcutaneous infusion of Ang II by pump implantation. Mice received Ang II for 3 or 28 days to mimic early and late stages of TAA development (Figures 2A). The effect of Ang II on hypertension was confirmed by the significant increase in systolic blood pressure measured after implantation, with no difference between Sdc-1^+/+^ and Sdc-1^-/-^ mice (data not shown). The use of this Ang II/BAPN model did not generate aneurysm in all studied animals (discussed in the following section). Analysis of Sdc-1 expression by IHC, revealed that Sdc-1 protein was elevated in TAA compared to healthy aortas after both 3 and 28 days of Ang II treatment (Figure 2B). Sdc-1 expression was specific for the TAA development as Sdc-1 was not detected in aortas treated only with BAPN or with both BAPN and Ang II (at both time points) and that did not develop aneurysm (Supplementary Figure 1). Therefore, in accordance with human sample data, Sdc-1 expression is increased in the TAA generated in this mouse model.

**Figure 2.**
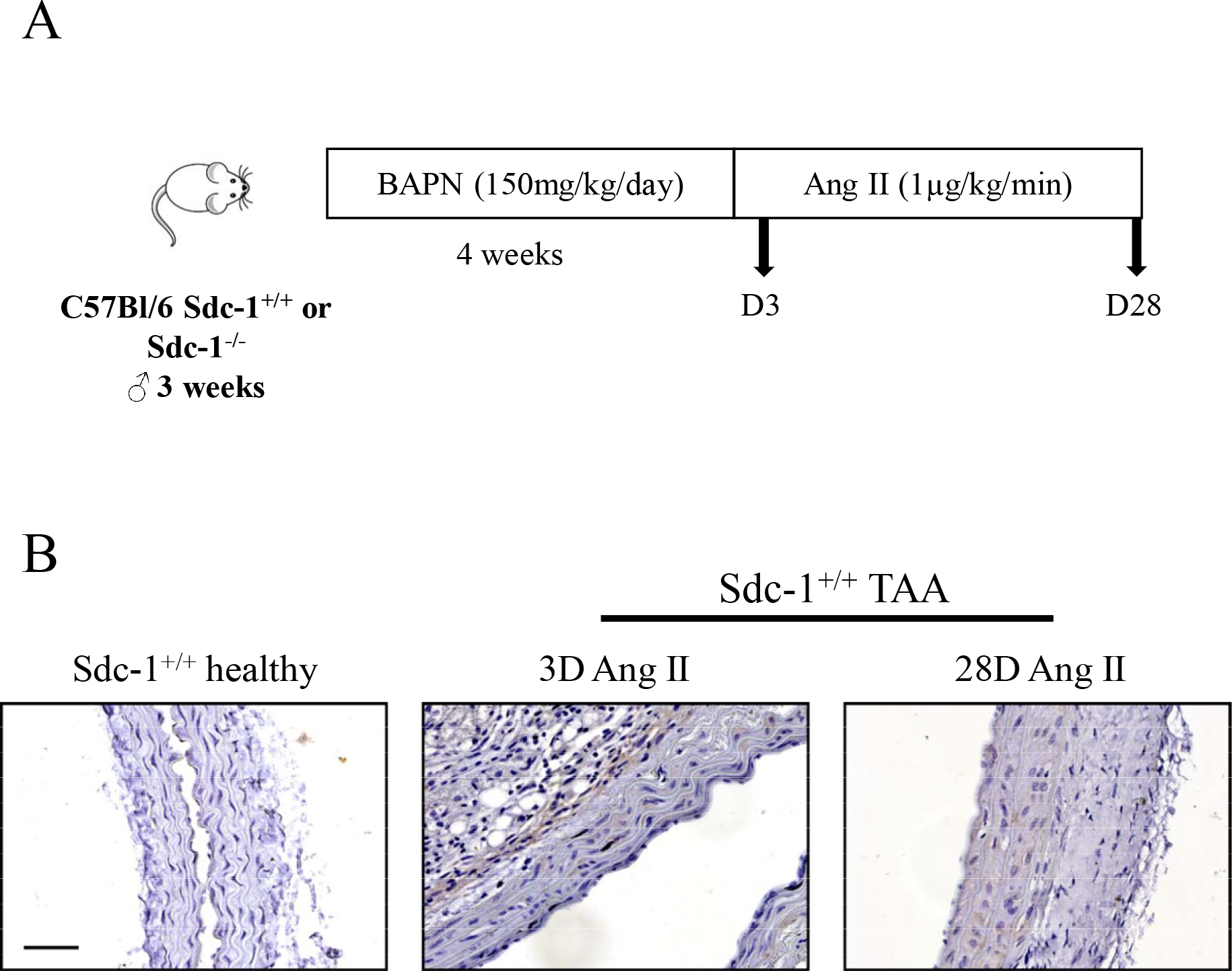
Sdc-1 is overexpressed in TAA compared to healthy aortas in the BAPN/AngII aneurysm mouse model. **(A)** Experimental model used. Sdc-1^+/+^ or Sdc-1^-/-^ male C57Bl/6J mice of 3 weeks old received intraperitoneal injection of β-amino propionitrile (BAPN) for 4 weeks followed by subcutaneous infusion of angiotensin II (Ang II) by pump implantation for 3 (D3) or 28 (D28) days, then sacrificed. **(B)** Representative images of Sdc-1 immunostaining in healthy (*n=3*) and TAA aortas after 3 (*n=2*) or 28 days (*n=3*) of Ang II treatment. Scale bar corresponds to 50µm.

### Sdc-1 is dispensable for TAA development and displays a potential protective role against AAA development in mice

The survival rate of Sdc-1^+/+^ and Sdc-1^-/-^ mice was similar during the course of Ang II infusion (Figures 3A), indicating that Sdc-1 deficiency did not alter the viability of the mice that developed aneurysm.

**Figure 3.**
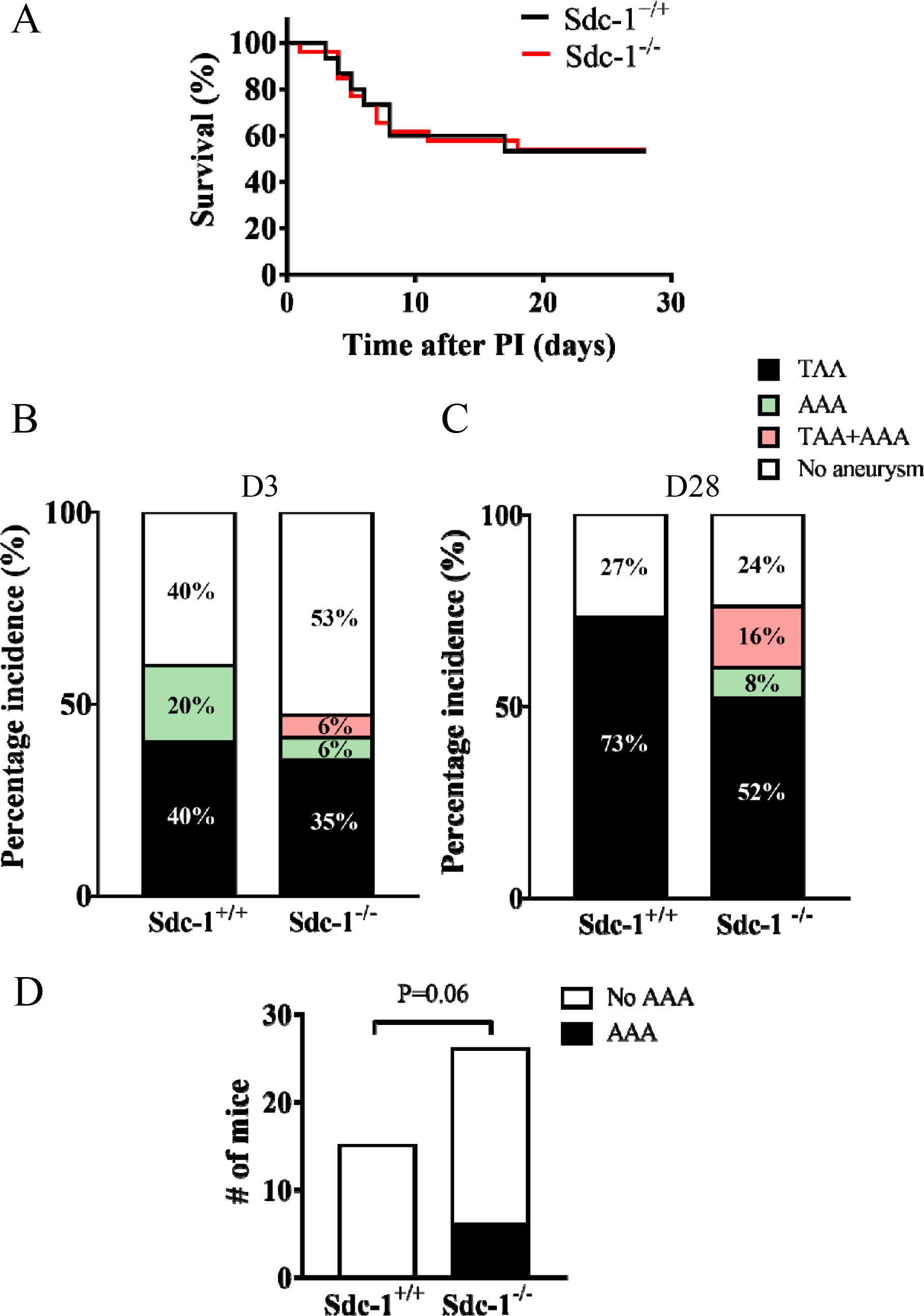
Sdc-1 is dispensable for TAA incidence or rupture but tends to protect from AAA development in mice. **(A)** Survival rate of mice after 28 days following Ang II infusion Gehan-Breslow-Wilcoxon test. PI : pump implantation. Sdc-1^+/+^ : *n=15*, Sdc-1^-/-^ : *n= 26*. ns, non-significant. **(B, C)** Percentage of TAA or AAA incidence in Sdc-1^+/+^ or Sdc-1^-/-^ mice for 3 **(B)** or 28 days **(C)** of Ang II treatment. (D) AAA incidence in mice treated with Ang II for 28 days. Fisher’s exact test. **(B)** Sdc-1^+/+^: *n=5*, Sdc-1^-/-^: *n= 17*. **(C, D)** Sdc-1^+/+^: *n=15*, Sdc-1^-/-^: *n= 26*.

As this model is known to induce both thoracic and abdominal aneurysms, TAA and AAA incidence were compared in Sdc-1^+/+^ and Sdc-1^-/-^ mice. After 3 days of Ang II infusion, the proportions of the developed aneurysms were as follow for Sdc-1^+/+^ and Sdc-1^-/-^ mice respectively: TAA (40% vs 35%), AAA (20% vs 6%), both TAA and AAA (0% vs 6%) (Figure 3B). These results did not reveal any significant implication of Sdc-1 in aneurysm incidence in this mouse model after 3 days of Ang II treatment.

When analyzing aneurysm incidence after 28 days of Ang II treatment, 73% of Sdc-1^+/+^ mice developed TAA vs 52% in Sdc-1^-/-^ mice. Moreover, 8% of Sdc-1^-/-^ developed AAA whereas none of Sdc-1+/+ mice did (Figure 3C). Sdc-1^-/-^ mice showed a higher tendency for AAA occurrence, alone or in combination with TAA compared to Sdc-1^+/+^ mice, with 6 /26 AAA in Sdc-1^-/-^ mice compared to 0/15 AAA in Sdc-1^+/+^ mice (Figure 3D). These data are in accordance with a previous study reporting a protective effect of Sdc-1 against AAA development in a mice model of AAA (22).

### Sdc-1 deficiency does not alter the extent of aortic dilatation, ECM remodeling, or leukocytes recruitment in descending TAA in mice

The observation that Sdc-1 was not involved in TAA incidence does not rule out the possibility that it could affect the extent of thoracic aortic dilatation or the morphology of the developed TAA.

Aortas harvested at 3 or 28 days from surviving animals were photographed (Figure 4A) and their external diameters were measured. There was no difference in the descending thoracic diameter between Sdc-1^+/+^ and Sdc-1^-/-^ aortas at both time points (Figure 4B). More specifically, we did not observe any difference in diameter between Sdc-1^+/+^ and Sdc-1^-/-^ aortas that displayed TAA, 28 days after pump implantation (data not shown).

**Figure 4.**
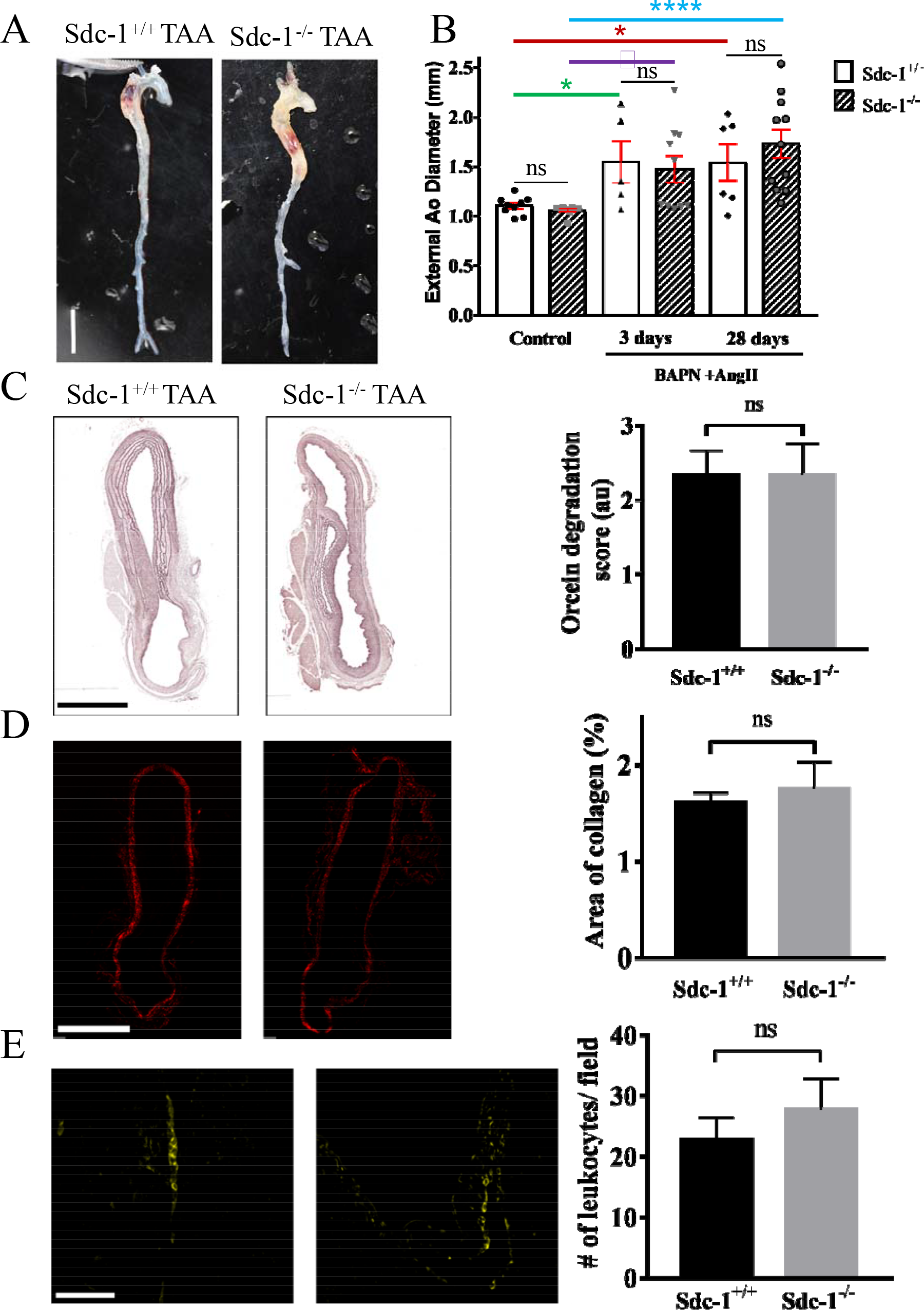
Sdc-1 deficiency does not alter the extent of aortic dilatation, ECM remodeling, or leukocyte recruitment in descending TAA in mice. **(A)** Macroscopic images of developed TAA in Sdc-1^+/+^ or Sdc-1^-/-^ mice. Scale bar corresponds to 5 mm. **(B)** Measurement of external descending thoracic diameter in Sdc-1^+/+^ and Sdc-1^-/-^ mice that received BAPN and Ang II for 3 or 28 days, and control (sham) mice. Aorta with a diameter ≥ 1.5 mm was considered as an aorta that developed TAA. Sdc-1^+/+^ or Sdc-1^-/-^ ctrl: *n= 9* for both, Sdc-1^+/+^ and Sdc-1^-/-^ 3 days: *n=5* and *n=10* respectively, Sdc-1^+/+^ and Sdc-1^-/-^ 28 days: *n=6* and *n=11* respectively. **(C)** Elastin degradation. Representative images of orcein staining for Sdc-1^+/+^ or Sdc-1^-/-^ TAA aortas after 28 days following Ang II treatment. Histological sections were evaluated for elastin degradation by giving an approximate score (scale from 0 to 4: 0 corresponding to no degradation and 4 to maximal degradation). Sdc-1^+/+^: *n=3*, Sdc-1^-/-^: *n= 6* **(D)** Collagen deposition. Representative images of Sirius red staining (visualized under polarized light) and its quantification by image J software. *P* values were calculated using two-tailed Mann-Whitney test. **(C, D)** Scale bar corresponds to 500µm. **(E)** Representative images of immunofluorescence staining of leukocytes (CD45) in TAA aortas after 28 days of Ang II infusion and its quantification (to the right) by Image J software. Scale bar corresponds to 50µm. **(D, E)** Sdc-1^+/+^: *n=3*, Sdc-1^-/-^: *n= 7*. **(B, C, D, E)** Data are presented as mean ± SEM and *P* values were calculated using two-tailed Mann-Whitney test;*: p<0.05, □: p<0.01,****: p<0.0001, ns: not significant.

The ECM remodeling of the developed Sdc-1^+/+^ or Sdc-1^-/-^ TAA was investigated by assessing the level of elastic degradation and collagen deposition by orcein and Sirius red staining, respectively. Similar levels of elastin degradation (Figure 4C) or collagen deposition (Figure 4D) were observed for Sdc-1^+/+^ and Sdc-1^-/-^ aortas with TAA.

Increasing evidence support a role of inflammation or immune cells infiltration in human TAA (reviewed in (24)) and TAA mice models are often associated with inflammation (25,26). In addition, Sdc-1 is involved in various aspects of inflammation such as leukocyte recruitment (27) or its resolution (28). Therefore, leukocyte and more specifically neutrophil recruitment were analyzed in Sdc-1^+/+^ or Sdc-1^-/-^ TAA aortas after 3 (Supplementary Figure 2A) or 28 days (Figure 4E, Supplementary Figure 2B) of Ang II infusion.

Massive recruitment of leukocytes (CD45 staining) and neutrophils (Ly-6G staining) in TAA was observed in a similar manner in both Sdc-1^+/+^ and Sdc-1^-/-^ mice following 3 days of Ang II infusion (Supplementary Figure 2A). Leukocytes and neutrophils were distributed nearly all over the aorta but sparsely in the intact media. These cells were mainly observed in the adventitia or at the border of the false channel and to a less extent in the dissected media part (media around false channel) (Supplementary Figure 2A). Less leukocyte infiltration was detected after 28 days of Ang II treatment compared to 3 days, with still no difference between Sdc-1^+/+^ and Sdc-1^-/-^ mice (Figure 4E, Supplementary Figure 2B). Moreover, very few neutrophils were observed in the TAA formed in both Sdc-1^+/+^ and Sdc-1^-/-^ mice at this time point (Supplementary Figure 2B).

Taken together, these results indicate that Sdc-1 has no effect on the extent of aortic dilatation, elastin degradation nor collagen deposition in descending TAA in mice. In addition, Sdc-1 does not participate in leukocytes (and neutrophils) recruitment in descending TAA in this mouse model.

## DISCUSSION

Previous studies suggested that Sdc-1 could be an important player in TAA development as it is involved in ECM assembly and organization (18), SMCs phenotype regulation and mechanosensing (19). RNA level of Sdc-1 (referred previously as syndecan) was previously reported to be increased after vascular injury (29). More recently, an increase of Sdc-1 protein level was observed in the adventitia of human aortas with ascending TAA (30).

In the current study, we report for the first time, by ELISA and histological analysis, that Sdc-1 is one of the members of the PGs overexpressed in human TAA aorta medial layer. Our immunofluorescence analysis showed that Sdc-1 expression is increased not only in media layer, but also in the intima and adventitia layers, in agreement with Ntika et al (30).

Our results also indicate that SMCs are the major cell type expressing Sdc-1 in human TAA aortas, which is in line with an *in vitro* study showing that mechanical stress induces the expression of Sdc-1 by SMCs (31). This suggests that the altered wall mechanics in TAA aortas induce Sdc-1 expression on SMCs.BAPN and Ang II infusion induce vascular remodeling (32), TAA, AAA (23), and thoracic aortic dissection (33). Our model was adapted from a study showing that BAPN and Ang II administration in C57Bl/6J mice without any specific genetic background, develop 49% of AAA and 38% of TAA mostly in the ascending part of the aorta (23). However, our Sdc-1^+/+^ C57Bl/6J mice did not develop any AAA and almost all of the TAA were found in the descending aorta, at the typical site of B dissection and not distal as found by others (23). The observed differences between these results can be explained by the different age of the mice at the beginning of the experiment (8 vs. 3 weeks), the route of BAPN administration (subcutaneous vs. intraperitoneal), the duration of Ang II treatment (6 vs. 4 weeks), or the mouse background (Sdc1^+/+^ from heterozygous mating vs commercially available mice).

Our results indicate that Sdc-1 is not involved in the incidence of descending TAA neither in the risk of rupture in our model. However, Sdc-1 deficiency has been reported to exacerbate AAA formation and rupture vulnerability in two mice models of abdominal aneurysms (22). We observe similar findings as AAA was observed only in Sdc-1^-/-^ mice and not in the Sdc-1^+/+^ mice, 28 days after Ang II infusion. The fact that Sdc-1 was not involved in the extent of aortic dilatation, elastin degradation or collagen deposition in TAA in our model suggests that Sdc-1 may have a more potent effect on ECM remodeling in abdominal rather than in thoracic aorta given their different structural and mechanical characteristics (34,35). The differences in the number of lamellar units, elastin and collagen content, proteinase system, and tension forces contribute to distinctive vascular remodeling at thoracic or abdominal sites (reviewed in (35)). Moreover, the inflammatory response is amplified in AAA compared to TAA (36), increasing the protease activity and MMPs production. It is worth to mention that PGs distribution is heterogeneous throughout the aorta providing a mechanism for regional dependent adaptation to variable hemodynamic stresses (37). More interestingly, the regulation of PGs has been shown to be distinct in the two types of the disease. For instance, aggrecan and versican accumulate in human ascending TAA (9), whereas proteomic analysis of AAA samples showed a reduced abundance of these PGs in comparison to healthy samples (38). Therefore, it would not be surprising that a single PG performs a distinct function specific to the site its of expression.

Notably, PGs display differential expression depending on the age of the individual (in mice or humans) and the severity of the disease (9,39–41). Examining the regulation of Sdc-1 in early human stages of TAA development is likely unachievable. We assumed that 3 days and 28 days of Ang II treatment corresponded to early and late stage of aneurysm, respectively. However, rupture (late stage of aneurysm) was observed all along the experimental protocol, even 24h after Ang II infusion. Thus, the used model did not permit us to study a possible contribution of this PG in early stages of the disease.

As observed for the human TAA samples, Sdc-1 protein expression increased in mice TAA compared to healthy aortas at both time points. The observed over expression in the media and adventitial layers of TAA aortas 3 days after Ang II treatment should correspond to both SMC and leukocyte expression. Indeed 28 days after Ang II infusion, Sdc-1 overexpression was observed only in the media and not the adventitia of TAA aortas, in concordance with the observation of decreased leukocytes infiltration at this time point.

We observed an infiltration of neutrophils localized mainly in the adventitia, in borders of the false channel and in dissected media in TAA aortas 3 days after Ang II infusion most likely because Ang II as a vasopressor, is a potent stimulant of neutrophils recruitment (42,43). Neutrophil infiltration was previously reported to be in intima following 24h of Ang II infusion (44). Our data showed that after a longer time of Ang II infusion (3 days), which corresponds to a more advanced stage of aneurysm, neutrophil infiltration took place in the media where dissection was observed and in the adventitia. However, these neutrophils almost disappeared 28 days after Ang II treatment, corresponding possibly to a switch from an acute to a chronic inflammatory phase.

Despite the findings that neutrophil Sdc-1 reduces neutrophil adhesion to the endothelium (15) and that Sdc-1 mediates neutrophils resolution by chemokines clearance (28), our data showed that Sdc-1 does not play a role in neutrophil recruitment in our model, since no difference was observed between Sdc-1^+/+^ or Sdc-1^-/-^ TA aortas. More investigations are still required to identify the identity of PGs present in the distinct types of thoracic aneurysm. The impact of pooled PGs/GAGs on medial degeneration could be either a global effect generated by all accumulated PGs, or specific where each PG by itself exerts a particular role during TAA development. A process of compensation (or redundancy) displayed by other PG expression in the present model could explain the absence of Sdc-1 specific effect, as it has been reported in the context of atherosclerosis between biglycan and perlecan for Apo-B retention (45).

## Conclusion

This study reports an overexpression of Sdc-1 in human TAA compared to healthy aortas suggesting that it can serve as a biomarker for this pathology. The underlying mechanism relevant to this alteration remains to be determined. The mouse model used induced mostly descending TAA, and Sdc-1 was not involved in its incidence, neither in its histological characteristics. An implication of Sdc-1 in TAA (ascending or descending) could be revealed from studies in other TAA mouse models possibly associated with ECM genes deficiency as it organizes the ECM and interacts with its proteins. Interestingly, we observed that Sdc-1 tends to protect from AAA, in agreement with a previous report (22). Deciphering the protective molecular function of Sdc-1 in AAA, and its possible role in TAA, could perhaps aid in the comprehension of its clinical relevance.

## Supporting information

Supplementary material Sdc-1 and Thoracic aneurysm

## Ethics Statement

Studies involving human participants were in agreement with the principles defined in the Declaration of Helsinki. Written informed consents were obtained from all individual participants (or family members) included in the study with Ethics Committee approval from INSERM and AP-HP (CEERB du GHU Nord) Institutional review board (CPP 05 04 32, Ambroise Paré, Boulogne, France, April 2005; updated in March 2008). Mice experiments were conducted according to the institutional guidelines for the care and use of animal research and the ARRIVE guidelines and in compliance with the Animal Care and Use Committee (2011-14/69-0035). The protocol was approved by the French MESRI (Ministère de l’Enseignement Supérieur de la Recherche et de l’Innovation # 8855)

## Author Contributions

BR conceived and designed the study; SZ performed most of the experiments; BR, SV and SJ performed some of human and/or mice experiments; MCB, VA, EH, YB, BR and SZ analysed the data; SZ wrote the manuscript with support from BR; MCB, VA, YB, OO, NC, EH, and BB supervised the research and provided intellectual discussion and editorial advice. All authors contributed to manuscript revision and approved the submitted version.

## Conflict of Interest

The authors declare that the research was conducted in the absence of any commercial or financial relationships that could be construed as a potential conflict of interest.

## Acknowledgments

We thank Thierry Dubois and Benoit Ho-Tin-Noé for the critical reading of the manuscript and helpful advices.

## Funding

This work was funded by INSERM and Sorbonne Paris Nord University. Sara Zalghout was funded by the Lebanese University, AZM Saade association, INSERM and Sorbonne Paris Nord University.

## Notes

### Competing Interest Statement

The authors have declared no competing interest.

