## Supplementary material Sdc-1 and Thoracic aneurysm for "Syndecan-1 is overexpressed in human thoracic aneurysm but is dispensable for the disease progression in vivo"

### Slide 1
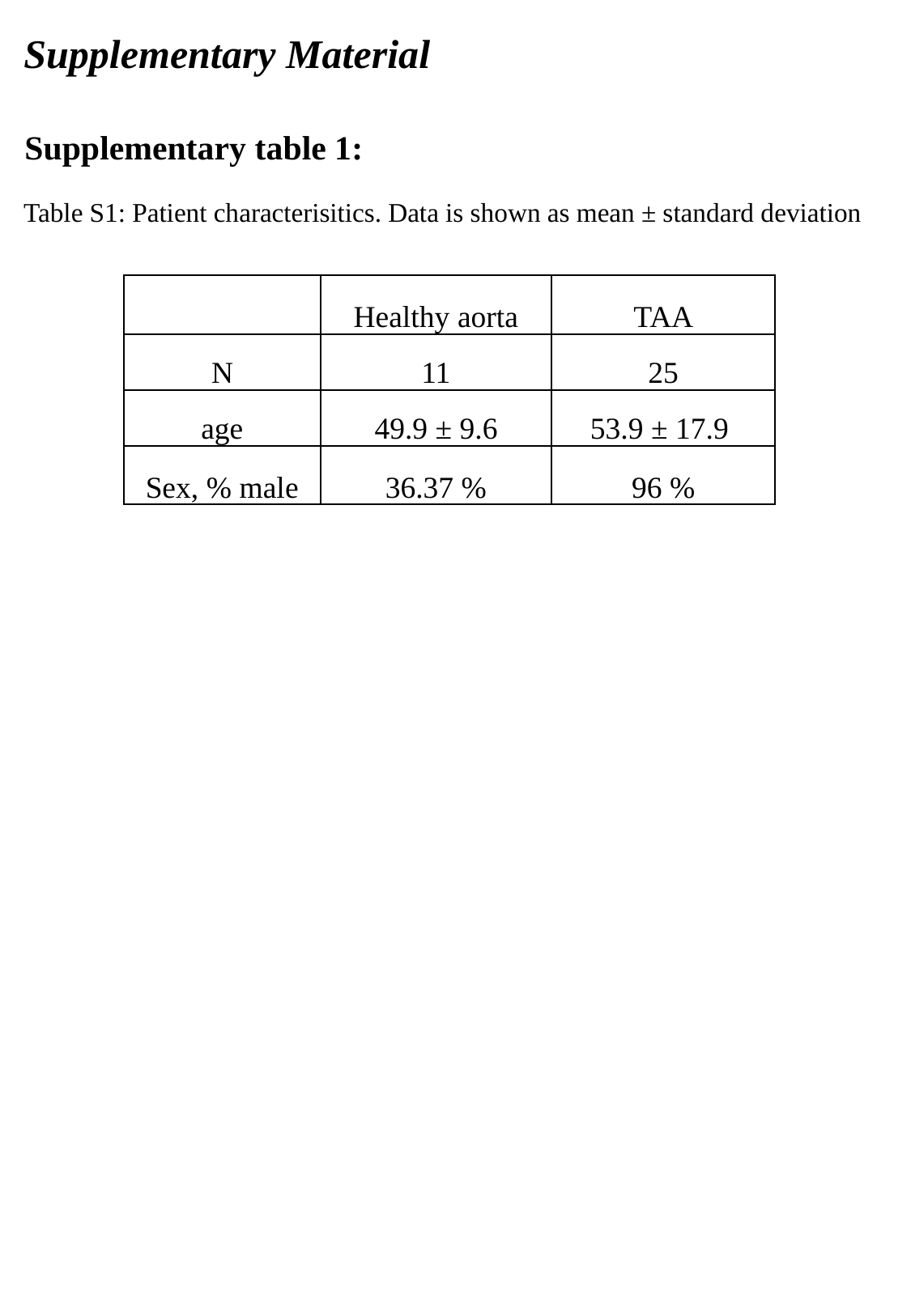

Supplementary Material
Supplementary table 1:
Table S1: Patient characterisitics. Data is shown as mean ± standard deviation
| | Healthy aorta | TAA |
| --- | --- | --- |
| N | 11 | 25 |
| age | 49.9 ± 9.6 | 53.9 ± 17.9 |
| Sex, % male | 36.37 % | 96 % |

### Slide 2
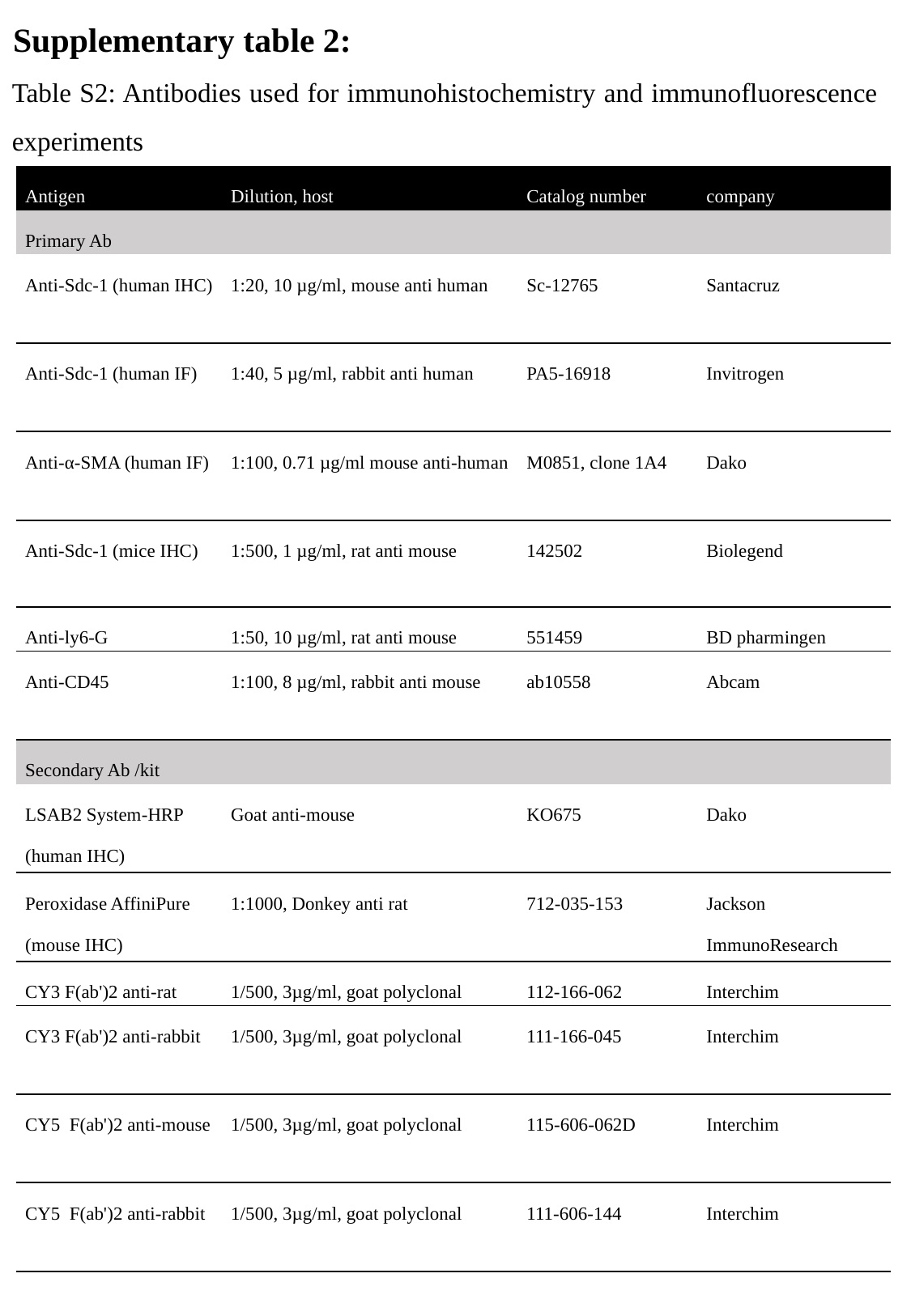

Supplementary table 2:
Table S2: Antibodies used for immunohistochemistry and immunofluorescence experiments
| Antigen | Dilution, host | Catalog number | company |
| --- | --- | --- | --- |
| Primary Ab | | | |
| Anti-Sdc-1 (human IHC) | 1:20, 10 µg/ml, mouse anti human | Sc-12765 | Santacruz |
| Anti-Sdc-1 (human IF) | 1:40, 5 µg/ml, rabbit anti human | PA5-16918 | Invitrogen |
| Anti-α-SMA (human IF) | 1:100, 0.71 µg/ml mouse anti-human | M0851, clone 1A4 | Dako |
| Anti-Sdc-1 (mice IHC) | 1:500, 1 µg/ml, rat anti mouse | 142502 | Biolegend |
| Anti-ly6-G | 1:50, 10 µg/ml, rat anti mouse | 551459 | BD pharmingen |
| Anti-CD45 | 1:100, 8 µg/ml, rabbit anti mouse | ab10558 | Abcam |
| Secondary Ab /kit | | | |
| LSAB2 System-HRP (human IHC) | Goat anti-mouse | KO675 | Dako |
| Peroxidase AffiniPure (mouse IHC) | 1:1000, Donkey anti rat | 712-035-153 | Jackson ImmunoResearch |
| CY3 F(ab')2 anti-rat | 1/500, 3µg/ml, goat polyclonal | 112-166-062 | Interchim |
| CY3 F(ab')2 anti-rabbit | 1/500, 3µg/ml, goat polyclonal | 111-166-045 | Interchim |
| CY5 F(ab')2 anti-mouse | 1/500, 3µg/ml, goat polyclonal | 115-606-062D | Interchim |
| CY5 F(ab')2 anti-rabbit | 1/500, 3µg/ml, goat polyclonal | 111-606-144 | Interchim |

### Slide 3
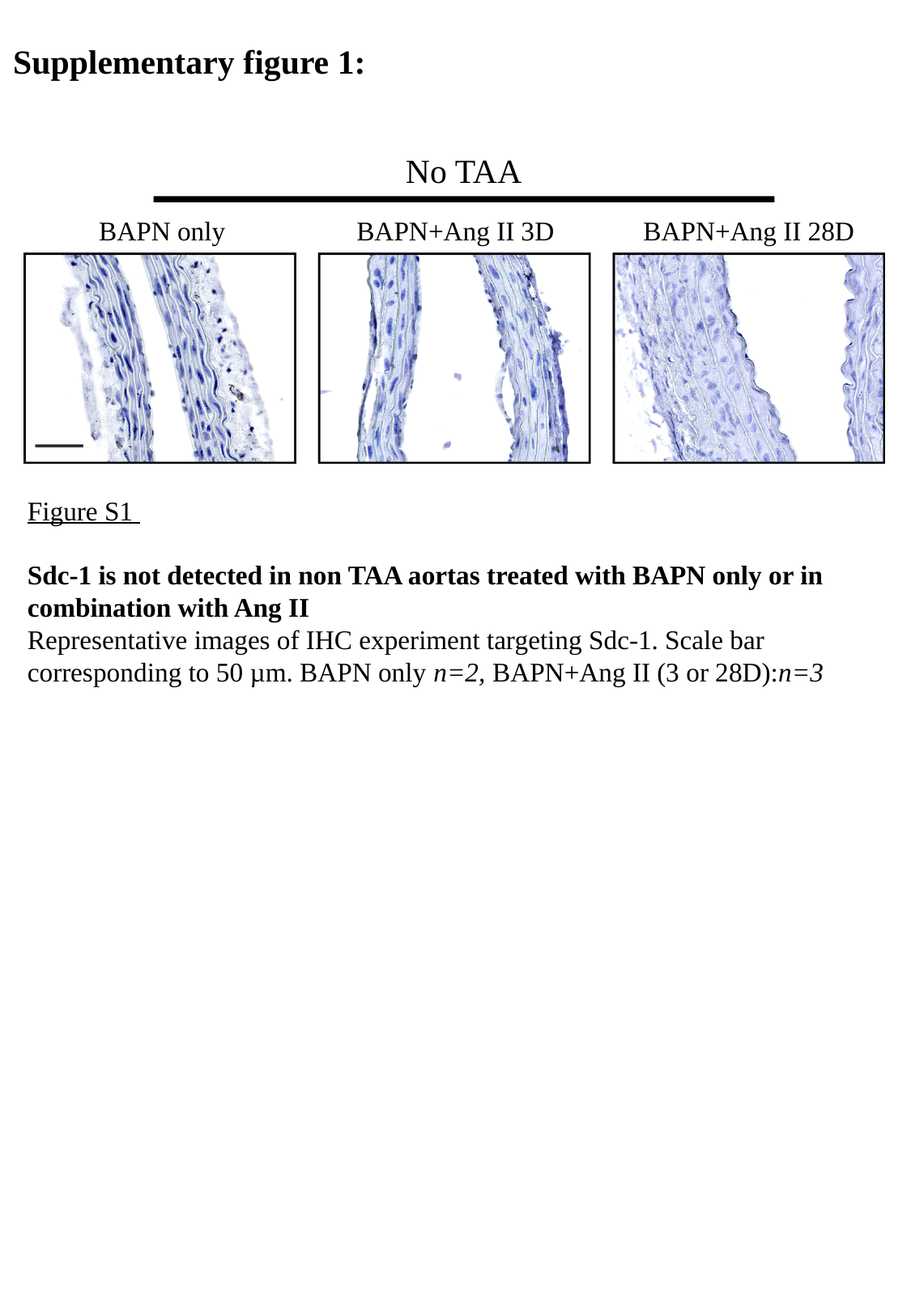

Supplementary figure 1:
No TAA
BAPN only
BAPN+Ang II 3D
BAPN+Ang II 28D
Figure S1
Sdc-1 is not detected in non TAA aortas treated with BAPN only or in combination with Ang II
Representative images of IHC experiment targeting Sdc-1. Scale bar corresponding to 50 µm. BAPN only n=2, BAPN+Ang II (3 or 28D):n=3

### Slide 4
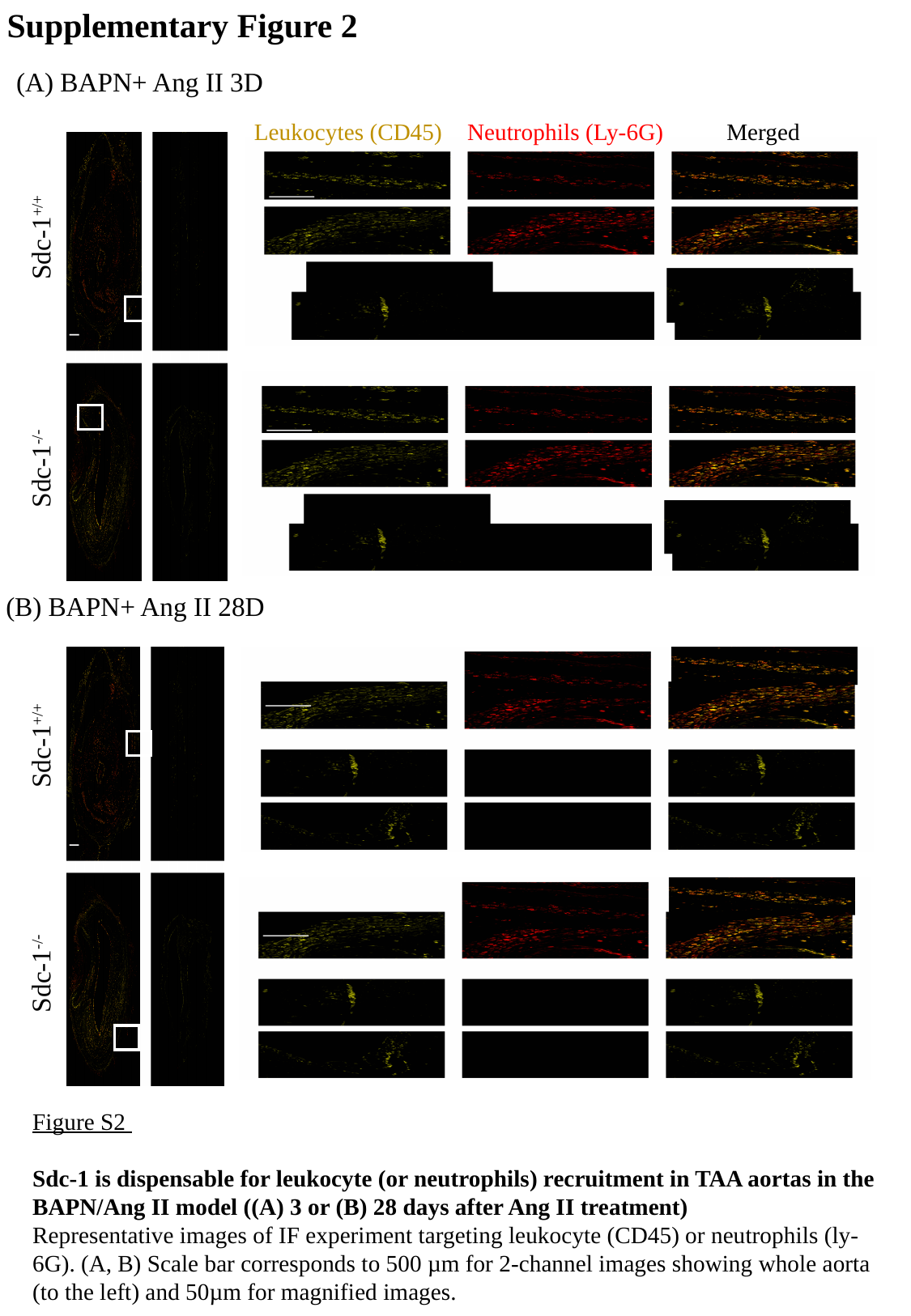

Supplementary Figure 2
(A) BAPN+ Ang II 3D
Merged
Neutrophils (Ly-6G)
Leukocytes (CD45)
Sdc-1^+/+^
Sdc-1^-/-^
(B) BAPN+ Ang II 28D
Sdc-1^+/+^
Sdc-1^-/-^
Figure S2
Sdc-1 is dispensable for leukocyte (or neutrophils) recruitment in TAA aortas in the BAPN/Ang II model ((A) 3 or (B) 28 days after Ang II treatment)
Representative images of IF experiment targeting leukocyte (CD45) or neutrophils (ly-6G). (A, B) Scale bar corresponds to 500 µm for 2-channel images showing whole aorta (to the left) and 50µm for magnified images.
